# DeepLUAD: An efficient approach for lung adenocarcinoma pattern classification

**DOI:** 10.1101/2022.05.06.490977

**Authors:** Ahmed Bouziane, Ala Eddine Boudemia, Taib Abderaouf Bourega, Mahdjoub Hamdi

**Author notes:** Corresponding author. (A. Bouziane). E-mail addresses* (A. E. Boudemia), (T. A. Bourega), (M. Hamdi).

## Abstract

Histopathological analysis of whole-slide images is the gold standard technique for diagnosis of lung cancer and classifying it into types and subtypes by specialized pathologists. This labor-based approach is time and effort consuming, which led to development of automatic approaches to assist in reducing the time and effort. Deep learning is a supervised classification approach that is well adapted for automatic classification of histopathological images. We aimed to develop a deep learning-based approach for lung adenocarcinoma pattern classification and generalize the proposed approach to the classification of the major non-small cell lung cancer types. Three publicly available datasets were used in this study. A deep learning approach for histopathological image analysis using convolutional neural networks was developed and incorporated into automatic pipelines to accurately classify the predominant patterns on the whole-slide images level and non-small cell lung cancer types on patch-level. The models were evaluated using the confusion matrix to perform an error analysis and the classification report to compute F1-score, recall and precision. As results, the three models have shown an excellent performance with best combination of hyper-parameters for training models. First and second models predicted adenocarcinoma predominant patterns on two different datasets with an accuracy, respectively, of 96.15% and 89.51%. The third model has exceeded an accuracy of 99.72% in classifying major non-small cell lung cancer types. The proposed deep learning-based lung cancer classification approach can be used to assist pathologists in identifying of lung adenocarcinomas patterns.

## Introduction

Lung cancer is the leading cause of cancer deaths worldwide, it is responsible for 1.8 million yearly deaths [1]. Lung cancer is grouped into two categories, non-small cell lung cancer (NSCLC) occupying 80% to 85% of lung cancer cases, and small cell lung cancer (SCLC), which represents the rest [2]. NSCLC is subdivided into three types: lung adenocarcinoma (LUAD), squamous cell carcinoma (SqCC), and most rarely, large-cell carcinoma (LCC). LUAD is the most frequent type among these three [3], and its treatment is determined by the tumor’s grade, stage, and subtype. The World Health Organization (WHO) categorizes LUAD into six subtypes: acinar, lepidic, cribriform, papillary, micropapillary, and solid, and suggests that tumors be identified according to predominant subtypes [4].

To categorize a patient’s diagnostised lung cancer, the pathologist inspect a huge amount of data (slides) using a microscope. This task is repetitive, very complex, and require hours of work for experienced pathologist. Furthermore, predictions vary from a pathologist to other due to many factors [5].

Many groups started developing computer-aided systems based on artificial intelligence (AI) techniques to help in the diagnosis of lung cancer, but most of them were restricted into classifying its major types (LUAD vs. SqCC) [6, 7]. However, accurate classification of LUAD growth patterns (subtypes) is often very challenging. Nearly 80% of LUAD cases are a mixed pattern of two or more histologic subtypes, and the classification is variable among pathologists since it depends on observation. [8]. Even though the observation of a biopsy by pathologists is considered by far the golden diagnostic procedure, but the accuracy of the assessment is lower than 80% [9].

The introduction of deep learning (DL) based approach is expected to improve classification accuracy of pathologists as it can reduce investigation time, reduce variability in interpretations and prevent overlooking. A DL algorithm exploits several layers of nonlinear information (transformations) [10] and is able to learn complex features by combining simple features already extracted from the data [11].

One approach is particularly interesting; it aims at extracting from each whole-slide image (WSI) a number of mini image patches. Since any typical WSI is at the megapixel level, extracting small patches (e.g.: 300*300 pixels) is very useful [12]. Furthermore, it allows to have a huge dataset of patches even from a small dataset of WSIs which helps with the training process, applying this method would facilitate feature extraction of tumors, help with the detection of even small tumor regions.

DeepPATH [13] is a deep convolutional neural network (CNN) with inception v3 architecture that trained on images from the cancer genome atlas (TCGA) in order to accurately classify WSIs into three classes: LUAD, SqCC or normal tissue. DeepPATH has shown a performance that matches pathologists with an average area under the curve (AUC) of 0.97.

DeepSlide is another more complex work [14]. By using a residual network (ResNet) model Deepslide can automatically classify the histologic patterns of LUAD and can reach pathologist level in this task. The model obtained a kappa score of 0.525 and had 66.6% agreement with three pathologists even higher than the inter-pathologist kappa score and agreement on this test set.

Gertych, A. et al [15] have proposed a pipeline that includes a CNN model recognize five classes: acinar, solid, cribriform, micropapillary, and non-tumor areas in LUAD. The best CNN achieved F1-scores of 0.91 (solid), 0.76 (micropapillary), 0.74 (acinar), 0.6 (cribriform), and 0.96 (non-tumor) respectively, with an accuracy of 89.24%.

Many researchers took advantage of the CNN to build systems that are able to diagnose the lung cancer, the most common challenge is the lack of publicly accessible, well-annotated data. Therefore, only few were able to develop systems, which classify the growth patterns of cancer or the subtypes and the performances of these models were less than 90%.

This paper proposes an approach called DeepLuad which greatly improves the classification accuracy of the five most frequent LUAD patterns using modern techniques of DL. It adopts small patch extraction, patches filtering, data augmentation, selection of the best parameters for the CNN model, patch-level predictions and WSI-level predictions. Furthermore, we have shown that our approach is generalizable to the classification of the major lung cancer types. The paper is divided into four sections: section 1 is an introduction. Section 2 describes materials and methods. The results are presented in section 3. Finally, section 4 draws discussion.

## 1. Materials and Methods

We mainly focused on training our model to classify LUAD subtypes on the first two datasets and then trained the model on a third dataset that contains major lung cancer types (LUAD vs SqCC vs Non Cancer tissues). An overview of the proposed approach is presented in Fig. 1.

**Fig. 1.**
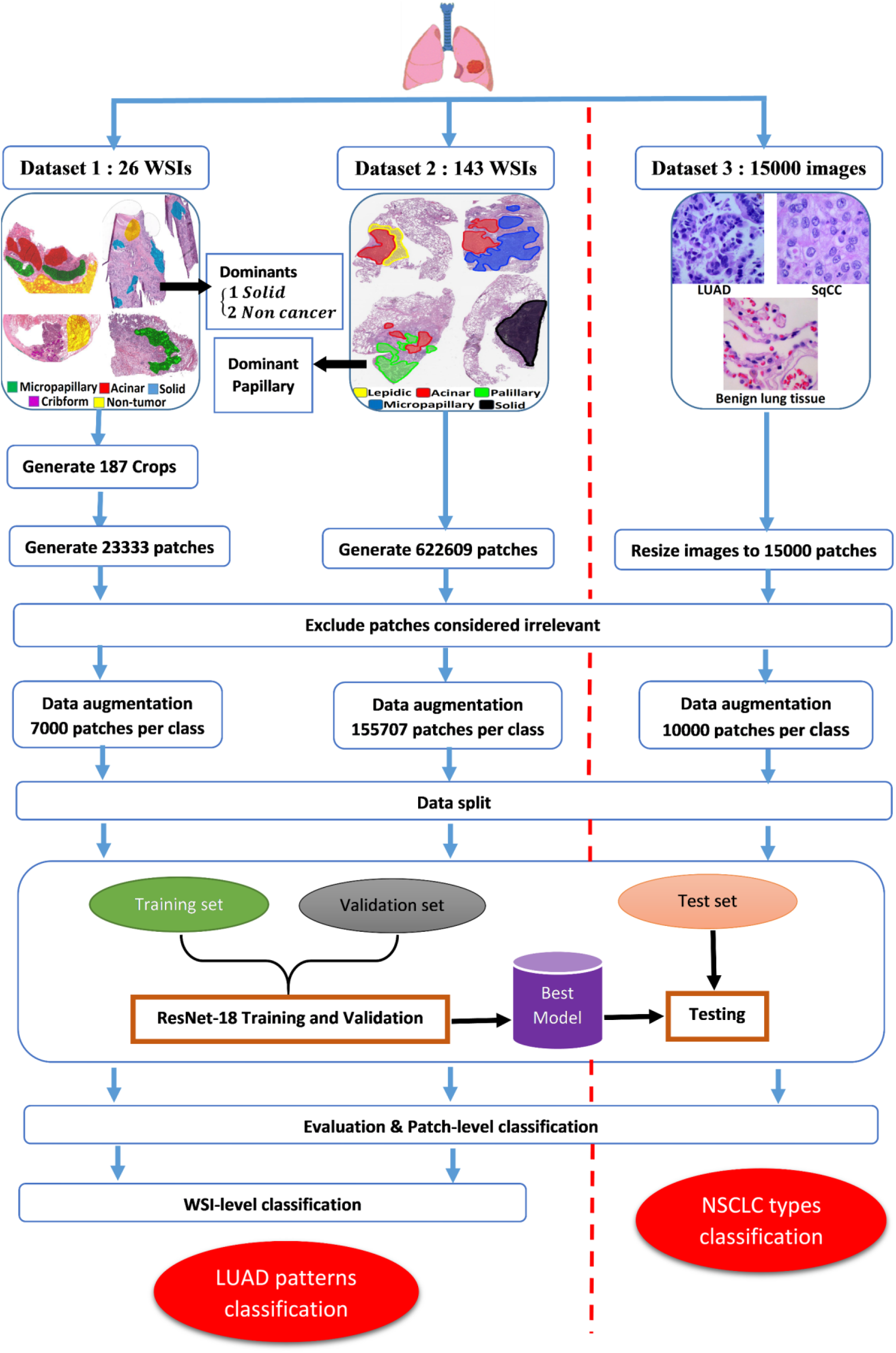
Overview of the proposed approach.

### 1.1. Dataset

Neural network usually requires a huge dataset that contains millions of training examples, but for medical image datasets this is a very common problem because large datasets are rare and even access to them is limited. Furthermore, even if we had access to a huge dataset such as the TCGA-LUAD, usually it would be poorly annotated and will not be of much use to train for some specific problems, such as LUAD subtypes classification where each image contains a mix of patterns. Bounding box annotations are required in this case.

In this work, we used digital histopathology images from three publicly available datasets. The first dataset consists of 26 WSIs from TCGA portal [16]. Cedars-Sinai Medical Center (CSMC) pathologists offered ground truth annotations of four subtypes of LUAD and non-tumor regions, these annotations are publicly available [15]. Pathologists examined the slides at 10x or greater magnification to detect tumor regions then lowered the magnification to circle and label tumor sites as solid, micropapillary, acinar or cribriform subtypes. Non-tumor regions were defined as those formed exclusively of non-cancerous (NC) components. Examples of LUAD subtypes are shown in Fig. 2

**Fig. 2.**
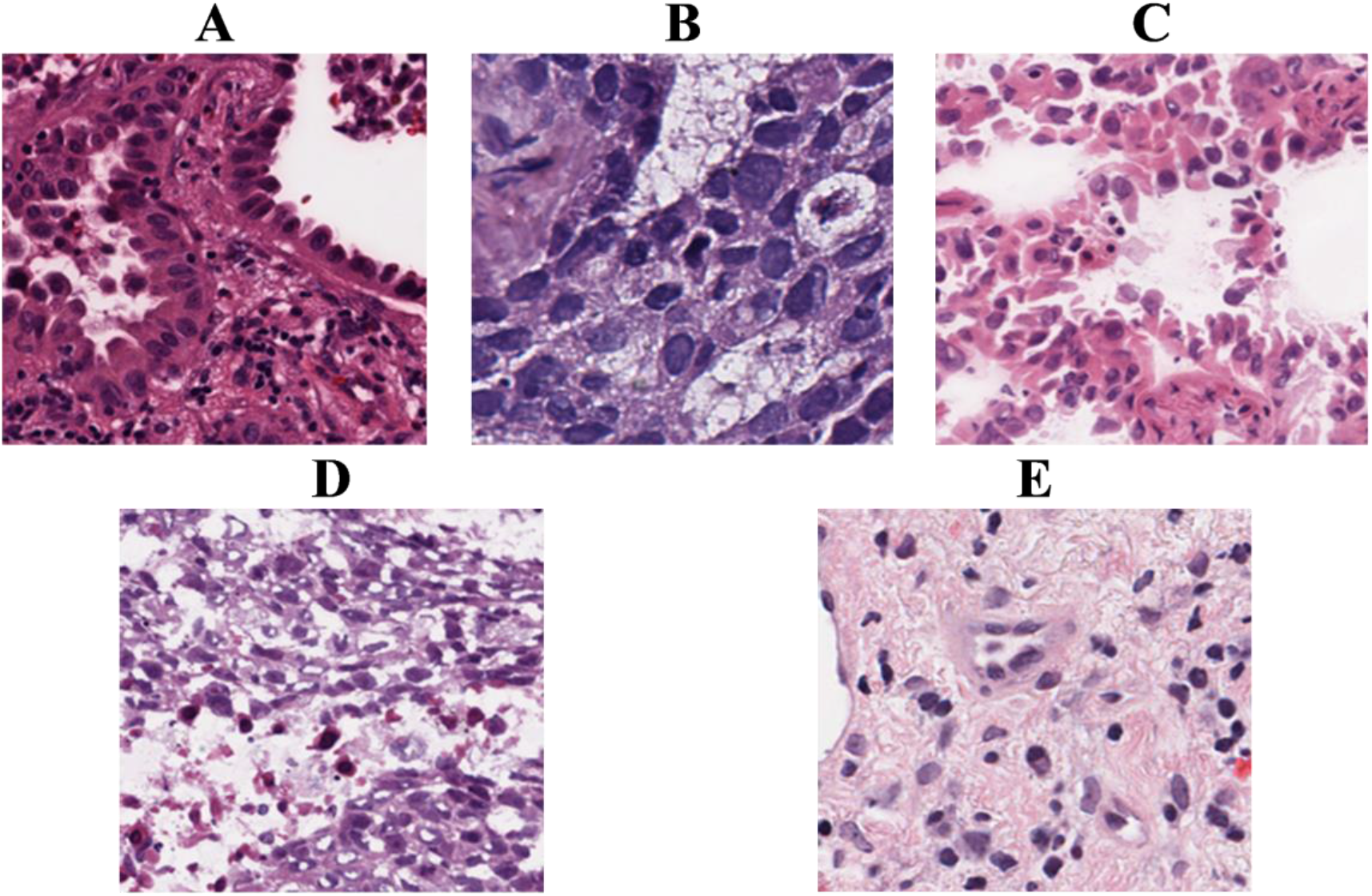
Examples of LUAD patterns and non-tumor regions from the first dataset. A) acinar, B) cribform, C) micropapillary, D) solid and E) NC [16].

The second dataset consists of 143 WSIs were obtained by Jason W. Wei. Et al [14]. The digital slides in this dataset were scanned at 20x magnification using a Leica Aperio whole-slide scanner at the Dartmouth-Hitchcock Medical Center (DHMC). Three pathologists from DHMC’s Department of Pathology manually labeled all WSIs. One or more of the five histological subtypes of LUAD can be found in each WSI (lepidic, solid, papillary, micropapillary and acinar). All images were labeled by the three pathologists, identifying the predominant subtypes on the WSI level. Dataset and annotations used in this study are available after a simple request procedure [17].

The third dataset has 750 images of lung tissue (250 benign lung tissue, 250 LUAD, and 250 SqCC) from pathology glass slides obtained by Andrew Borkowski et al [18]. The original authors augmented the images using the augmentor software tool [18], and the final dataset was increased to 15,000 RGB images divided into three classes. Furthermore, each class contains 5000 images. All images have a size of 768 x 768 pixels and saved in jpeg format. Dataset used in this study is publicly available [19].

### 1.2. Preprocessing

#### 1.2.1. First dataset

The first step was to generate crops of different sizes using the XML annotation files which have the coordinates of each crop and its label and then generate patches of size (448,448,3) that were compressed to (224,224,3). The reason for that is to get a bigger window sliding over the WSIs and covering a decent area that can be interpreted by the model.

We read crops from the corresponding WSIs and return it as an RGBA image (stands for Red-Green-Blue Alpha color space). We excluded irrelevant patches by eliminating any patch that is mostly white and saved only patches with decent purple area to be classified. Patches are then saved as TIFF files since it is the basis for SVS format and it is a lossless format to make sure we did not lose any important detail. We were able to generate a total number of 23333 patches from a total number of 187 crops. Since some classes were extremely underrepresented we had to use an overlapping factor different from one, therefore we changed the overlapping factor values. The distribution of histologic patterns is listed in Table 1.

**Table 1.** First dataset distribution for each subtype.

| Histologic pattern | Acinar | Cribriform | Micropapillary | Non-cancerous | Solid |
| --- | --- | --- | --- | --- | --- |
| **Crops** | 22 | 4 | 23 | 53 | 85 |
| **Patches (no-overlap)** | 1328 | 85 | 4053 | 6222 | 5207 |
| **Patches (overlap)** | 4507 (0.5) | 921 (0.3) | 5706 (0.75) | 6222 (1) | 5207 (1) |

#### 1.2.2. Second dataset

In the second dataset the labeling step was made by pathologists at the whole-image level. Therefore, instead of generating patches corresponding to each crop like the first dataset, we generated patches with the same label for each image. Which gives us less performance compared to the first dataset. For the preprocessing part of this dataset, the steps are the same as the first dataset. This time, we have not used the overlapping factor to generate patches, and the distribution of the histologic patterns is presented in Table 2.

**Table 2.**
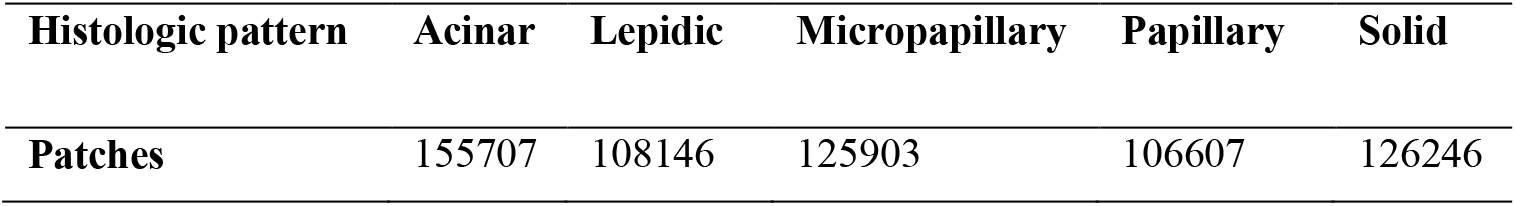
Second dataset distribution for each subtype.

| Histologic pattern | Acinar | Lepidic | Micropapillary | Papillary | Solid |
| --- | --- | --- | --- | --- | --- |
| Patches | 155707 | 108146 | 125903 | 106607 | 126246 |

#### 1.2.3. Third dataset

Since the patches are already generated and annotated, we only applied an algorithm to resize the images from (768, 768, 3) to (224, 224, 3) which is a better fit to the used model. We used the same procedure employed in the first dataset to eliminate irrelevant patches. We finally save the resized image as a JPEG image.

### 1.3. Data augmentation

The main advantages is to gain more data for training, helps a lot for balancing classes and it optimizes the process of learning relevant features for the model [20]. If a class were significantly over-represented, the model would penalize it the same way as other classes and, therefore, this class would contribute to the loss much more than the others. This will lead to a biased model that tends to classify any new image to the majority class increasing the rate of false positives and false negatives. To solve this problem, the implemented techniques in this work are: rotation [90, 180, 270], Gaussian blur with radius = 1, horizontal flip, vertical flip. Images are selected randomly; therefore, each image has the possibility to be augmented up to 15 times. For the first dataset, we augmented the number of images to 7000 per class. For the second dataset, we augmented the number of images to 155707 per class. For the third dataset, we performed an augmentation with a number of patches of 10000 per class.

### 1.4. Data splitting

We distributed the data into a train set, validation set and test set with the distribution ratios 80%, 10%, 10%, respectively, for the first dataset and 60%, 20%, 20%, respectively, for the second dataset. We randomly choose images and distribute them into validation and test sets while the rest will go to train set. We made sure that each augmented image will go to the same set as the original image; this is just a precaution to prevent the model from memorizing patterns instead of learning valuable features. For the third dataset, we used the same distribution ratios employed by Abbas, M. A. [21] to perform comparison, 55% for train, 20% for validation and 25% for test.

### 1.5. Deep neural network

Several studies have shown that the deeper a neural network goes the better its learning results especially for CNNs in image classification tasks. However, that goes to a certain point in CNNs that stack layer after layer in a traditional way, after that point the quality of the results have shown to be degrading with depth. However, this time the reason is not the famous phenomenon of overfitting but it is what we call vanishing gradient. ResNet was suggested to face this problem in deep CNNs by using residual mapping of stacked layers.

When the network is too deep, the gradients that help calculating the loss function drops to zero quickly. This means that no weights will be updated and therefore no training or learning is happening. Using residual blocs means that the gradients will pass directly from the last layers to initial filters via the shortcut connections. ResNet uses a shortcut that approximates many nonlinear layers to zero for approaching identity identification. This does not make the network more complex or adds additional parameters but it solves the problem of vanishing gradients in deep networks. ResNet is also excellent complex identification tasks such as analyzing medical images [22]. The basic building block of ResNet is shown in Fig. 3.

**Fig. 3.**
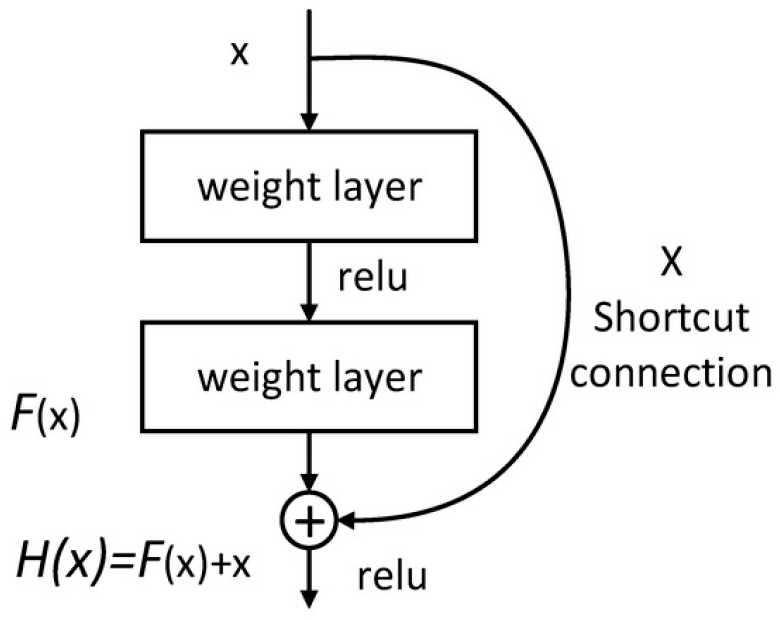
Basic structure of ResNet.

In this study, we choose to work with resnet-18 with a pre-trained model from Pytorch framework [23] that is designed to output (classify) 1000 classes, therefore, we had to modify the last part of the network to output only five classes corresponding to the different histologic patterns we are working on. We randomly initialized weights for the last layer that we modified.

Retraining those weights is called fine-tuning. Using a pre-trained model has the benefits of reducing training time, since the model does not need to learn low level features like edges, because these features are the same in almost every context and the model will focus only on learning high-level features that are specific to our data. Transfer learning is also very useful for problems where there are not so much data. In our case, generated patches are too precious to let the model learn basic features from them. The ResNet-18 was fine-tuned with multi-class cross-entropy as the loss function. Cross entropy can be employed for binary or multi-class classification where the target (image) contains one of the classes for example (cancer vs. non-cancer tissue) or (SqCC vs. LUAD vs. normal tissue). However, what is critical for us, it is also used in multi-label classification where one target can contain multiple classes together. In our case, this is helpful as most slides are contain a mix of LUAD subtypes [24]. From a logistical point of view, cross-entropy is particularly important, as it contributes to the estimation of models used in determining the probability of rare events, which are often the most expensive in terms of cost. This indicator differs significantly from other more straightforward precision indicators [25]. Furthermore, one of the most convincing arguments is that cross-entropy provides large gradient values, which are particularly interesting for the gradient descent that is the most efficient method of scale optimization at present [26].

There are many optimization algorithms but only few can be generalized over time, the most widely used one is gradient descent and its variants stochastic gradient descent, root mean square propagation (RMSprop), momentum and Adam. In this work we selected Adam, which is one of the newest and most efficient algorithms for gradient descent optimization. Adam calculates the exponential average of the gradient as well as the squares of the gradient for each parameter; the learning rate is then multiplied by the average of the gradient and by dividing it by the square root of the exponential average of the gradients, then the update is added [27].

### 1.6. Training, validation, and testing

We trained the network for 200 epochs for the first and the second datasets and 50 epochs for the third dataset. From experiment result, best results obtained by using learning rate value 1e-03 for LUAD pattern classification and 1e-06 for Major NSCLC types classification.

Training data was distributed into mini-batches of size 16, and we shuffled this data to assure that the batches contain different subclasses of cancer. We always save the best model by comparing the validation loss of the current batch with the lowest validation loss from the previous batch; this will ensure that we always save the model with the best weights.

After training and validation process, we ran the models on the test set to determine the final accuracy. The metrics that we used are the confusion matrix and the classification report. The confusion matrix allows us to perform an error analysis and see in which subclass the model is making more false predictions. The confusion matrix allows us to compute F1-score, recall and precision.

### 1.7. WSI-level prediction

This is only valid for the first and the second datasets. Since our model predicts in patches, we had to find a way to aggregate those predictions to determine the predominant subtype in each slide. We used the prediction file, performed data cleaning where we eliminated the predictions coming from augmented patches, and only kept those coming from original patches. We then grouped patches coming from the same WSI, averaged the probabilities of all valid predictions and looked for the subclass with the highest probability to predict the predominant pattern.

## 2. Results

While training a model it is very important to validate the results of each iteration. This is done by monitoring the validation loss compared to the training loss. The overall evolution of these two parameters should always tend to be minimized. Comparing Validation loss with training loss is not sufficient to validate a model; therefore, we needed to compute the accuracy of the model at the end of each iteration. The overall evolution should increase until hitting a plateau.

### 2.1. LUAD pattern classification

Results of the evaluation metrics for the first dataset are shown in Fig. 4. The first graph Fig. 4A compares the evolution of the validation loss with the training loss per epochs. It clearly shows that the both, training and validation loss, are decreasing steadily and following a normal behavior, which indicates that the model is learning/training properly, and the weights are being updated. Therefore, the model is getting better at predicting each class with every epoch. At the end, values were equal to 0.0029 for the training loss and 0.021 for the validation loss.

**Fig. 4.**
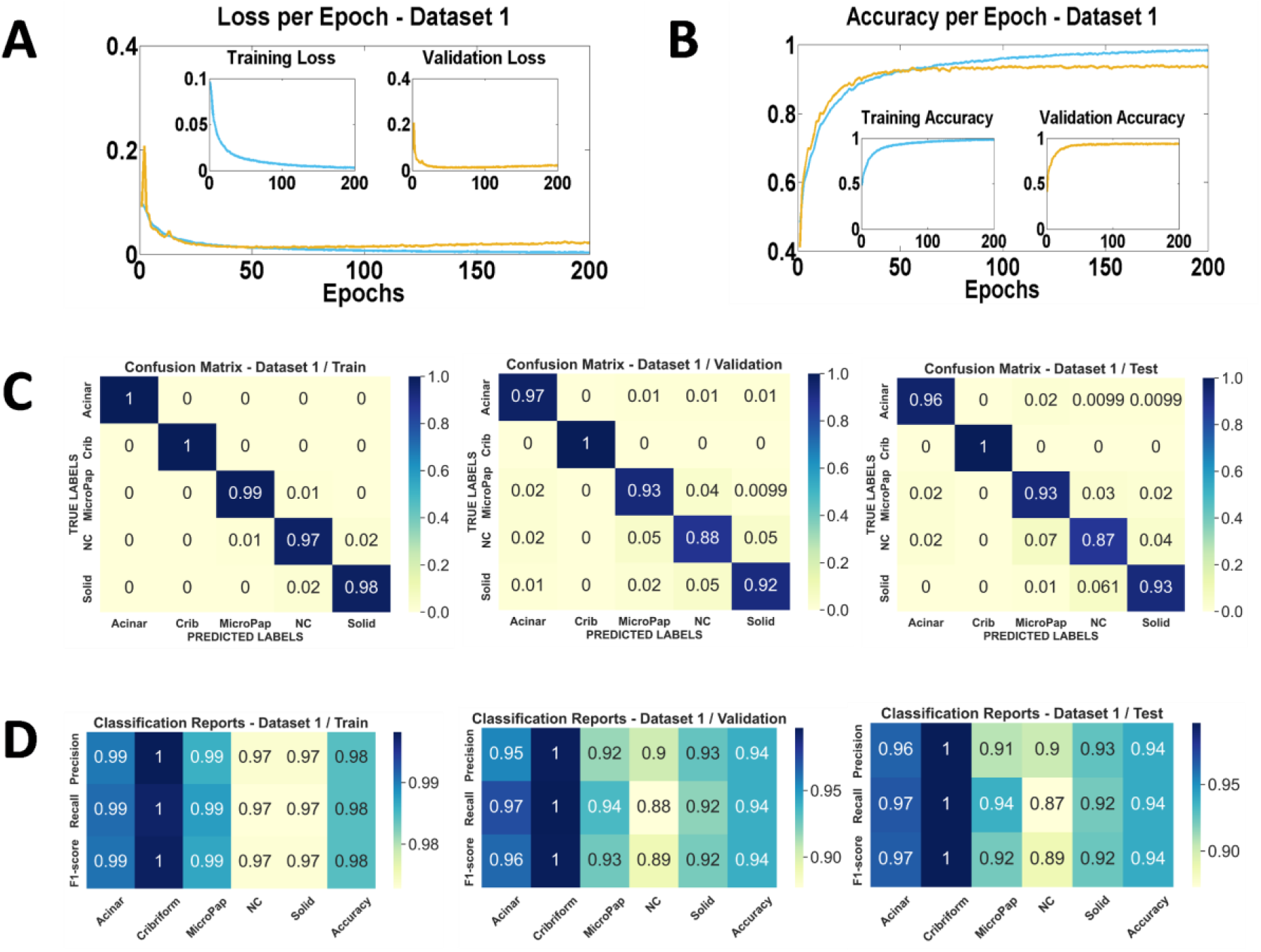
Results of the evaluation metrics for the first dataset. A) Loss per epoch, B) Accuracy per epoch, C) Confusion matrix of Train, Validation and Test (from left to right), D) Classification reports of Train, Validation and Test (from left to right).

The second graph in Fig. 4B shows the accuracy of both validation and training per epochs. We can immediately observe an upward trend for both curves and there is no significant difference at the beginning, except that the training accuracy went up to 98.46% and the validation one stabilized at 93.68%. From these two graphs, we can conclude that the model had a good behavior and was able to get astonishing results. For the patch-level predictions, when we ran our trained model on the testing set, we achieved an accuracy of 93.92%.

If we compare our results to the results of Gertych et al. [15], we can see that we achieved better performance with fewer resources, they were able to generate 797,680 image tiles for training, in our case we only generated 40,000. Still, our testing accuracy was far better than what they achieved as they only managed to get 84% while we have 93.92% for the accuracy of the patch-level predictions.

In each of the confusion matrices presented in Fig. 4C, we can see that most of the data were correctly classified since we can observe a diagonal line on the heat-map that indicates that the predicted labels and the correct ones are intersecting for the majority of images and the results were very satisfying. From the classification reports presented in Fig. 4D, we can conclude that the model is performing best on the Cribform class, followed by the Acinar class and almost in a similar way for the rest of the classes.

Results of the evaluation metrics for the second dataset are presented in the following Fig. 5. In the first graph presented in Fig. 5A, it can be clearly observed that the training loss curve is following a downward trend and decreasing steadily. At the same time, we can see some fluctuations in validation loss, especially in the first 50 epochs, which are due to the properties of the Adam optimizer. The fluctuations were disappeared in the second half of the graph, and the validation loss almost stable with a very interesting value around 0.02. Overall, the curves are indicating a great process of learning.

**Fig. 5.**
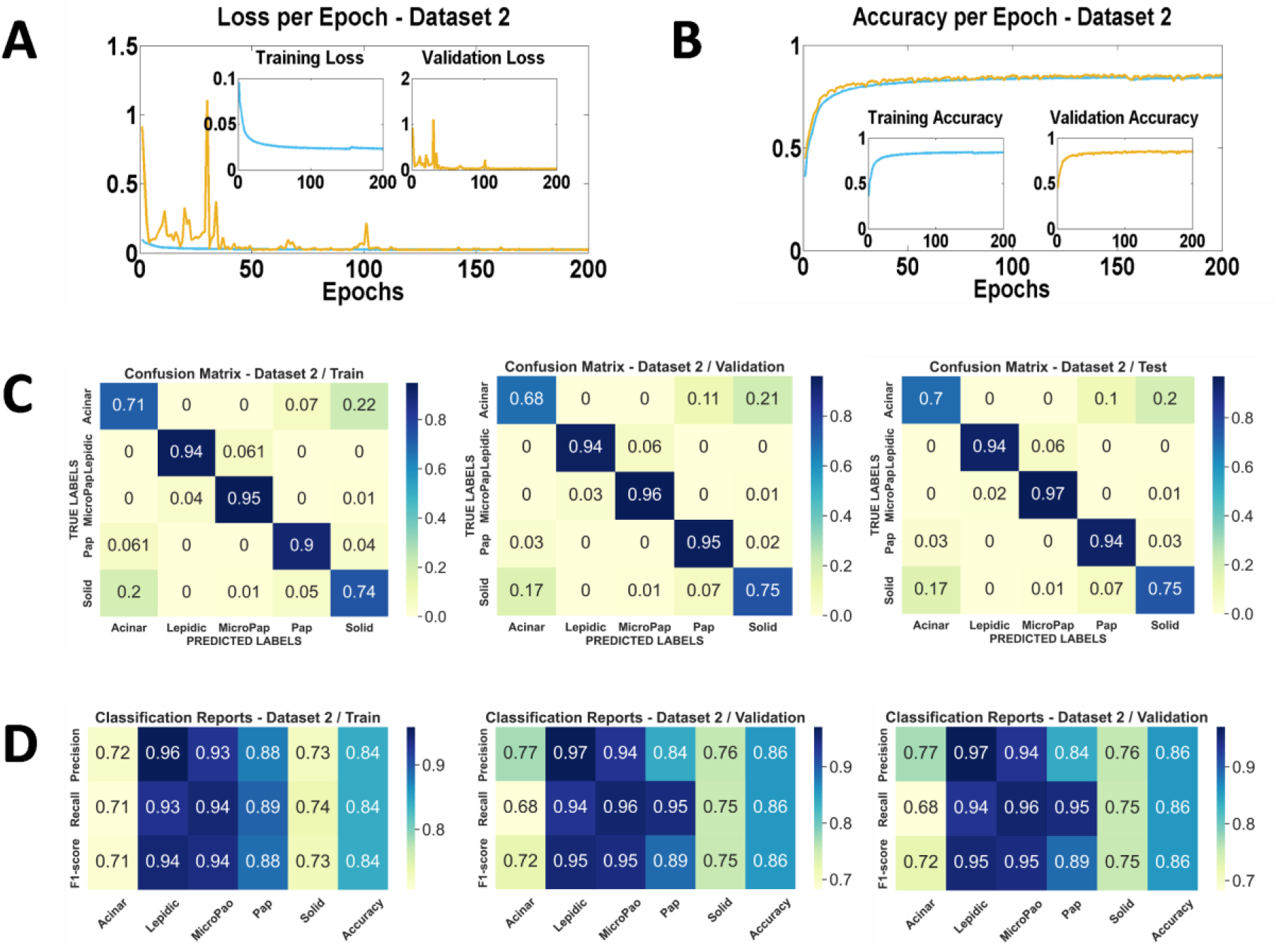
Results of the evaluation metrics for the second dataset. A) Loss per epoch, B) Accuracy per epoch, C) Confusion matrix of Train, Validation and Test (from left to right), D) Classification reports of Train, Validation and Test (from left to right).

We can immediately observe an upward trend for both curves in the accuracy graph (Fig. 5B). The model could learn quickly and stabilize at an accuracy of 86%. When we ran our trained model on the test set, we achieved an accuracy of 85.60%.

Despite the promising results found with this dataset, the decrease compared to the first dataset is very logical and due to the difference in annotation between the two datasets. For the first dataset, all the regions of each image are circled and labeled, on the other hand, for the second dataset the labeling is done for the whole image, so less detail with this last dataset, which will influence the final accuracy result. In addition, due to the large number of patches with WSI-level predictions for this second dataset, in the case where there are several cancer subtypes in the same image, unlike the first dataset, unfortunately the image is labeled only by the predominant subtype.

The confusion matrices and classification ratios shown in Fig. 5C-D, respectively, give very good results, with an apparent diagonal concentration in confusion matrices that proves the correct classification of the majority of patches, especially for the predominant subtypes.

According to the previous explanation, for the other misclassified patches, they belong to the other minority subtypes present in each image. This explanation is confirmed in the following paragraph by the increase of the accuracy value in WSI-level predictions. For metrics shown in the classification report, they have excellent scores with such difficulty.

For the WSI-level predictions, the following Table 3 shows a comparison between our results and the ground truth annotation of some examples. In the first dataset the obtained results were very satisfying and promising to predict the predominant subtype of the tissue with an accuracy of 96.15% and the rest 3.85% correspond only to one case, where there is a presence of two cancer subtypes plus non-tumorous components in the tissue. For this case, in the annotation provided from pathologists, non-tumorous components presents the predominant region followed by micropapillary and solid respectively, in our results micropapillary presents the predominant region followed by non-tumorous components and solid respectively.

**Table 3.**
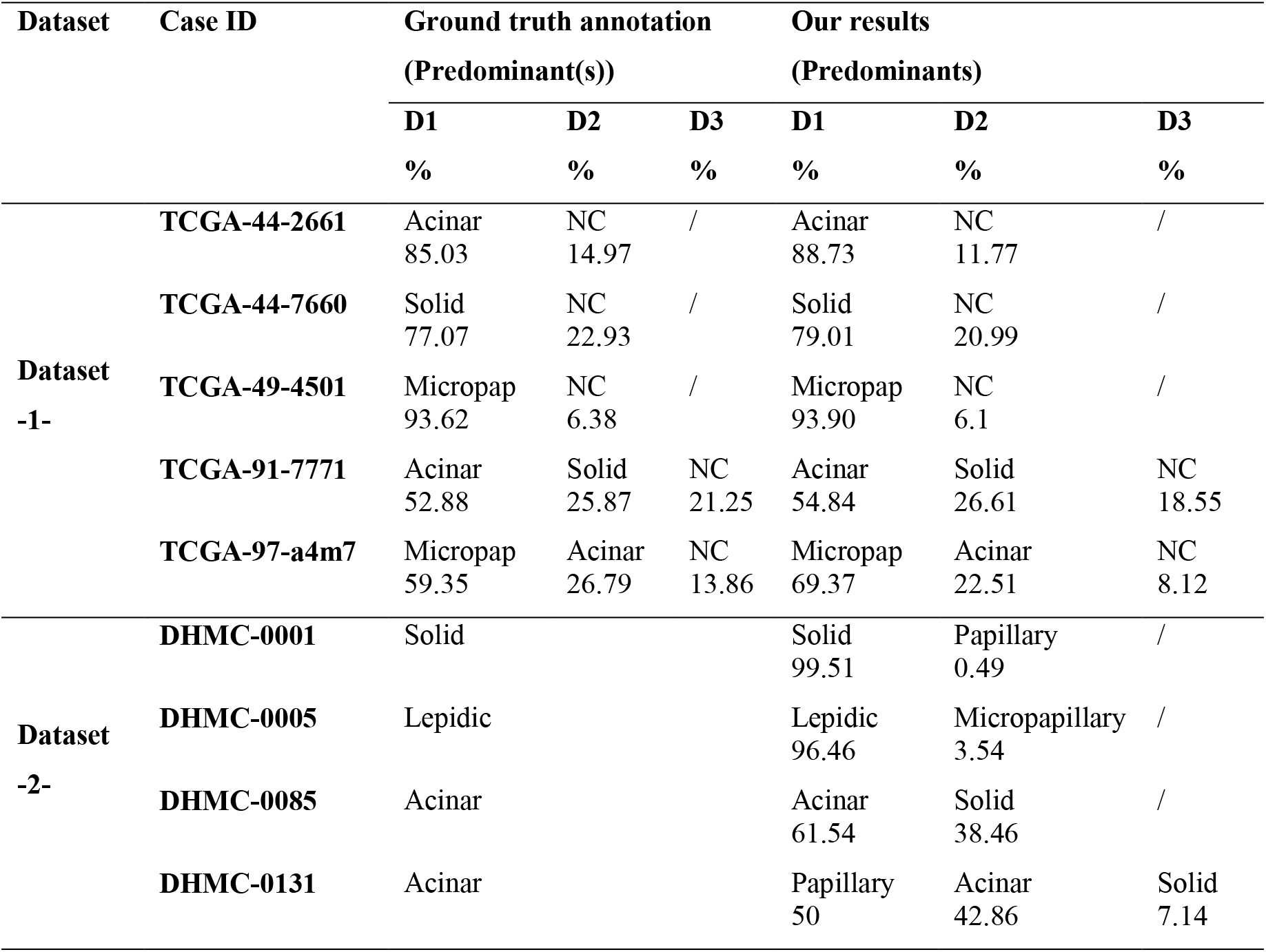
Whole slide level prediction examples for the first and second datasets.

| Dataset | Case ID | Ground truth annotation<br>(Predominant(s)) |  |  | Our results<br>(Predominants) |  |  |
| --- | --- | --- | --- | --- | --- | --- | --- |
|  |  | D1 | D2 | D3 | D1 | D2 | D3 |
|  |  | % | % | % | % | % | % |
| Dataset<br>-1- | TCGA-44-2661 | Acinar<br>85.03 | NC<br>14.97 | / | Acinar<br>88.73 | NC<br>11.77 | / |
|  | TCGA-44-7660 | Solid<br>77.07 | NC<br>22.93 | / | Solid<br>79.01 | NC<br>20.99 | / |
|  | TCGA-49-4501 | Micropap<br>93.62 | NC<br>6.38 | / | Micropap<br>93.90 | NC<br>6.1 | / |
|  | TCGA-91-7771 | Acinar<br>52.88 | Solid<br>25.87 | NC<br>21.25 | Acinar<br>54.84 | Solid<br>26.61 | NC<br>18.55 |
|  | TCGA-97-a4m7 | Micropap<br>59.35 | Acinar<br>26.79 | NC<br>13.86 | Micropap<br>69.37 | Acinar<br>22.51 | NC<br>8.12 |
| Dataset<br>-2- | DHMC-0001 | Solid |  |  | Solid<br>99.51 | Papillary<br>0.49 | / |
|  | DHMC-0005 | Lepidic |  |  | Lepidic<br>96.46 | Micropapillary<br>3.54 | / |
|  | DHMC-0085 | Acinar |  |  | Acinar<br>61.54 | Solid<br>38.46 | / |
|  | DHMC-0131 | Acinar |  |  | Papillary<br>50 | Acinar<br>42.86 | Solid<br>7.14 |

In the second dataset, given the nature of the dataset and the labeling on the WSI-level, specifying only the predominant pattern for each image, once again the obtained results were satisfying to predict the predominant subtype of the tissue with an accuracy of 89.51%. The rest 10.49% almost for all predominant subtypes labeled by pathologists, they were identified as second predominant by our model as for example the case ID DHMC-0131.

### 2.2. Major NSCLC types classification

For the third dataset and as predicted, the loss curves shown in Fig. 6A are following a downward trend. Both the losses decreased steadily with some fluctuations in validation loss until they stabilized at 0.0003 for the training loss and 0.0012 for the validation loss.

**Fig. 6.**
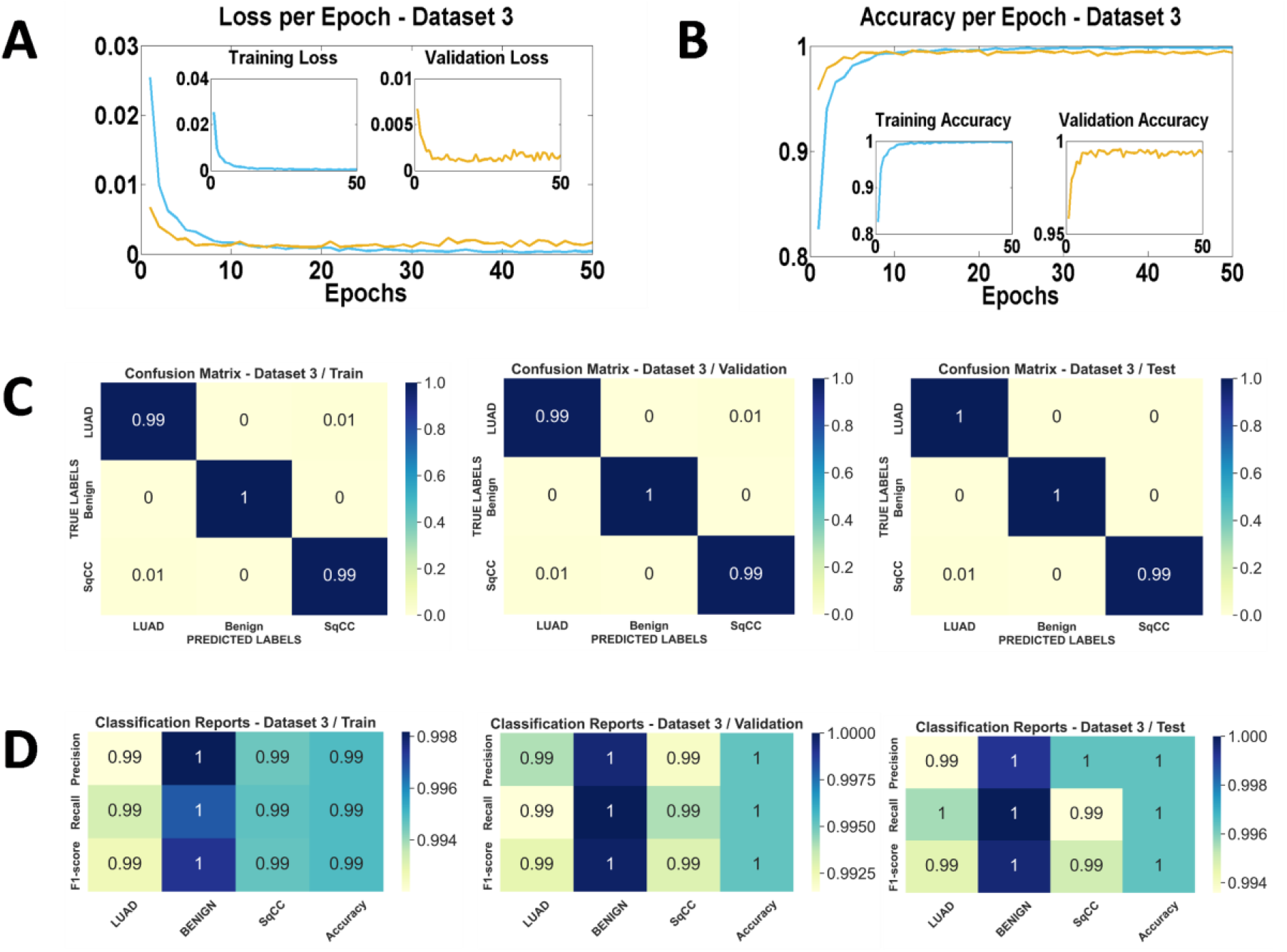
Results of the evaluation metrics for the third dataset. A) Loss per epoch, B) Accuracy per epoch, C) Confusion matrix of Train, Validation and Test (from left to right), D) Classification reports of Train, Validation and Test (from left to right).

For the accuracy graphs (Fig. 6B), there is an upward trend starting from 82.61% for the training and from 95.84% for the validation until finally achieving an accuracy of 99.89% for the training and 99.56% for the validation. One hypothesis is that a pre-trained model is already a model that performs well on low-level features and shapes and since cells were not very complicated structures, at least on a two-dimensional level, the model was able to have high accuracy from the beginning of the training phase. When we ran our trained model on the test set, we achieved an accuracy of 99.72%.

A comparative study was published by Abbas, et al. [21], where six state-of-the-art CNNs were used to perform the task we did with this third dataset. Comparative results are presented in Table 4.

**Table 4.** Comparative Table.

| CNN | Alex-Net | ResNet-18 | ResNet-34 | ResNet-50 | ResNet-101 | VGG 19 | Our Results |
| --- | --- | --- | --- | --- | --- | --- | --- |
| **Accuracy** | 97.26% | 98.8% | 99.18% | 99.6% | 99.8% | 98.93% | 99.72 |

In our approach using ResNet-18 we were able to achieve an accuracy of 99.72%, which is higher than five models in the study and very close to ResNet-101. But comparing them, our approach is the best choice, since ResNet-101 has approximately 42 million trainable parameters while in our approach using ResNet-18, we use only 11.17 million trainable parameters, which help to reduce the computational expense. This confirms that we achieved outstanding performance with our strategy.

The confusion matrices presented in Fig. 6C are almost perfect. With only so little wrong classifications compared to the total number of cases. The matrices show a great amount of correctly predicted data, forming a diagonal line on the heat map.

The results of the classification report confirm the previous analysis; the model has achieved great results and has high performance, as shown in Fig. 6D.

## 3. Discussion

Our approach has proven to be a very powerful method. This was statistically demonstrated by the evaluation metrics that we used. Our first model edged out pathologists who have an estimated assessment accuracy of 80% to 85% for the diagnosis of lung cancer patterns on histologic slides, for us we have found a test accuracy of 93.92% for the patch-level predictions and 96.15% for the WSI-level predictions. These results imply that we have gotten very close to the “Bayes Error,” considering that the majority of LUAD cases have mixed patterns in them. It can be challenging to outline the exact limits between two or more patterns. Finding and classifying minor patterns is even more challenging. This is probably because minor patterns can be easily overlooked or neglected, which leads to differences in annotations. Our model, on the other hand, is working with a “divide and conquer” strategy by processing each patch pixel by pixel before aggregating the results to predict which subtype is predominant in a particular slide. The effectiveness of our approach was confirmed by the results found in the second dataset. In the third dataset, we achieved almost perfect results with over 99% testing accuracy to classify the types of cancer which is not as hard as classifying its subtypes.

Our strategy has proven to be solid; we were able to find the best combination of hyper-parameters for training. Furthermore, our pipeline can be generalized and tested to any other medical imaging diagnosis task. Model used in the third dataset can easily be implemented with the model of the first and second datasets in a more general pipeline, where we determine the type of cancer before engaging in classifying its subtypes if it appears to be a LUAD.

Usually, researchers tend to normalize the images to reduce or augment the intensity of H&E coloration. They even modify the hue, contrast, and saturation of an image, trying to force the model to neglect these features, but we did not do that in our work, and this might be one of the reasons that our model achieved excellent performance. In fact, when we tried this kind of manipulation we did see a lot of over-fitting and we had to implement regularization methods to fight it back.

The study concludes that using such models in computer-aided diagnosis will help to reduce human error and make the whole process easier and more efficient. Our models are a perfect example of the efficiency of DL in multi-class classification. If implemented in laboratories, it will be a huge benefit to pathologists and researchers in lung cancer. However, while our pipeline can be used for this task, it can be tested on different types of cancer.

## CRediT authorship contribution statement

*Ahmed Bouziane*: Conceptualization, Methodology, Software, Validation, Formal analysis, Investigation, Resources, Data Curation, Writing - Original Draft, Writing - Review & Editing, Visualization, Supervision, Project administration.

*Ala Eddine BOUDEMIA*: Conceptualization, Methodology, Software, Validation, Formal analysis, Investigation, Data Curation, Writing - Original Draft.

*Taib Abderaouf Bourega*: Conceptualization, Methodology, Software, Validation, Formal analysis, Investigation, Data Curation, Writing - Original Draft.

*Mahdjoub Hamdi*: Conceptualization, Methodology, Software, Formal analysis, Investigation, Resources, Writing - Review & Editing, Visualization.

## Acknowledgments

This research is part of PRFU Project-D00L05ES250220220005 supported by the Algerian Directorate General for Scientific Research and Technology Development (DGRSDT-MESRS). The authors would like to thank Gertych, A et al, Wei, J.W. et al, and Borkowski, A. A et al for making the datasets publicly available. The authors would also like to thank the center of High Performance Computing (HPC) IbnKhaldoun of Biskra University in Biskra for the use of their computational resources.

